# Identifying and correcting repeat-calling errors in nanopore sequencing of telomeres

**DOI:** 10.1101/2022.01.11.475254

**Authors:** Kar-Tong Tan, Michael K. Slevin, Matthew Meyerson, Heng Li

**Affiliations:** Department of Medical Oncology, Dana-Farber Cancer Institute, Boston, MA, USA; Cancer Program, Broad Institute of MIT and Harvard, Cambridge, MA, USA; Department of Genetics, Harvard Medical School, Boston, MA, USA; Center for Cancer Genomics, Dana-Farber Cancer Institute, Boston, MA, USA; Department of Data Sciences, Dana-Farber Cancer Institute, Boston, MA, USA; Department of Biomedical Informatics, Harvard Medical School, Boston, MA, USA

**Keywords:** Nanopore-sequencing, long-reads, telomere, basecalling

## Abstract

Nanopore long-read genome sequencing is emerging as a potential approach for the study of genomes including long repetitive elements like telomeres. Here, we report extensive basecalling induced errors at telomere repeats across nanopore datasets, sequencing platforms, basecallers, and basecalling models. We found that telomeres which are represented by (TTAGGG)_n_ and (CCCTAA)_n_ repeats in many organisms were frequently miscalled (~40-50% of reads) as (TTAAAA)_n_, or as (CTTCTT)_n_ and (CCCTGG)_n_ repeats respectively in a strand-specific manner during nanopore sequencing. We showed that this miscalling is likely caused by the high similarity of current profiles between telomeric repeats and these repeat artefacts, leading to mis-assignment of electrical current profiles during basecalling. We further demonstrated that tuning of nanopore basecalling models, and selective application of the tuned models to telomeric reads led to improved recovery and analysis of telomeric regions, with little detected negative impact on basecalling of other genomic regions. Our study thus highlights the importance of verifying nanopore basecalls in long, repetitive, and poorly defined regions of the genome, and showcases how such artefacts in regions like telomeres can potentially be resolved by improvements in nanopore basecalling models.

## Background

Telomeres are protective caps found on chromosomal ends, and are known to play critical roles in a wide range of biological processes and human diseases [1,2]. These highly repetitive structures enable cells to deal with the “end-replication problem” through the action of telomerase which adds telomeric repeats to the ends of chromosomes. In cancer, the reactivation of telomerase to drive telomere elongation is estimated to occur in as many as 90% of human cancers, and has been shown experimentally to be critical for malignant transformation [3–8]. As one ages, telomeres are also known to progressively shorten, and are thus thought to also play a central role in the process of aging [9–11]. In many organisms, telomeres are characterized by (TTAGGG)_n_ repeats that vary in length of between 2 and 20kb long, which are not readily resolved by short-read sequencing approaches. Given the importance of telomeres in a wide range of biological process and the technical challenges associated with their analysis using short-read sequencing, there is significant interest in applying emerging techniques like long-read sequencing to study these repetitive structures.

Long-read sequencing has emerged as a powerful technology for the study of long repetitive elements in the genome. Two main platforms, Single Molecule Real Time (SMRT) sequencing, and Nanopore sequencing, have been developed to generate sequence reads of over 10 kilobases from DNA molecules [12,13]. In SMRT Sequencing, the incorporation of DNA nucleotides is captured real time via one of four different fluorescent dyes attached to each of the four DNA bases, thereby allowing the corresponding DNA sequence to be inferred. Sequencing of the same DNA molecule multiple times in a circular manner further allows highly accurate consensus sequence of the DNA molecule to be generated in a process termed Pacific Biosciences (PacBio) High-Fidelity (HiFi) sequencing [12]. During Nanopore sequencing, the ionic current, which varies according to the DNA sequence, is measured while a single-stranded DNA molecule passes through a nanopore channel. The electrical current measurement is then converted into the corresponding DNA sequence using a deep neural network trained on a collection of ionic current profiles of known DNA sequences [13]. Notably, both platforms enable long DNA molecules of more than 10 kilo-base-pairs to be routinely sequenced and are thus highly suited for the study of long repetitive elements like telomeres.

## Results and discussion

In our analysis of telomeric regions with nanopore long-read sequencing in the recently sequenced and assembled CHM13 sample [14,15], we surprisingly observed that telomeric regions were frequently miscalled as other types of repeats in a strand-specific manner. Specifically, although human telomeres are typically represented by (TTAGGG)_n_ repeats **(Supplementary Figure 1a)**, these regions were frequently recorded as (TTAAAA)_n_ repeats **(Figure 1a,b, Supplementary Figure 1 and 2a)**. At the same time, when examining the reverse complementary strand of the telomeres which are represented as (CCCTAA)_n_ repeats, we instead observed frequent substitution of these regions by (CTTCTT)_n_ and (CCCTGG)_n_ repeats **(Figure 1a,b, Supplementary Figure 1 and 2b,c)**. Notably, these artefacts were not observed on the CHM13 reference genome [14,15], or PacBio HiFi reads from the same site **(Figure 1a,b)**, suggesting that these observed repeats are artefacts of Nanopore sequencing or the base-calling process, rather than real biological variations of telomeres. Further, these repeat-calling errors could be observed on all chromosomal arms for the CHM13 sample **(Supplementary Figure 1b,c),** and were thus not restricted to a single chromosomal arm. The examination of each telomeric long-read also indicates that these error repeats frequently co-occur with telomeric repeats at the ends of each read **(Figure 1c, Supplementary Figure 3)**.

**Figure 1.**
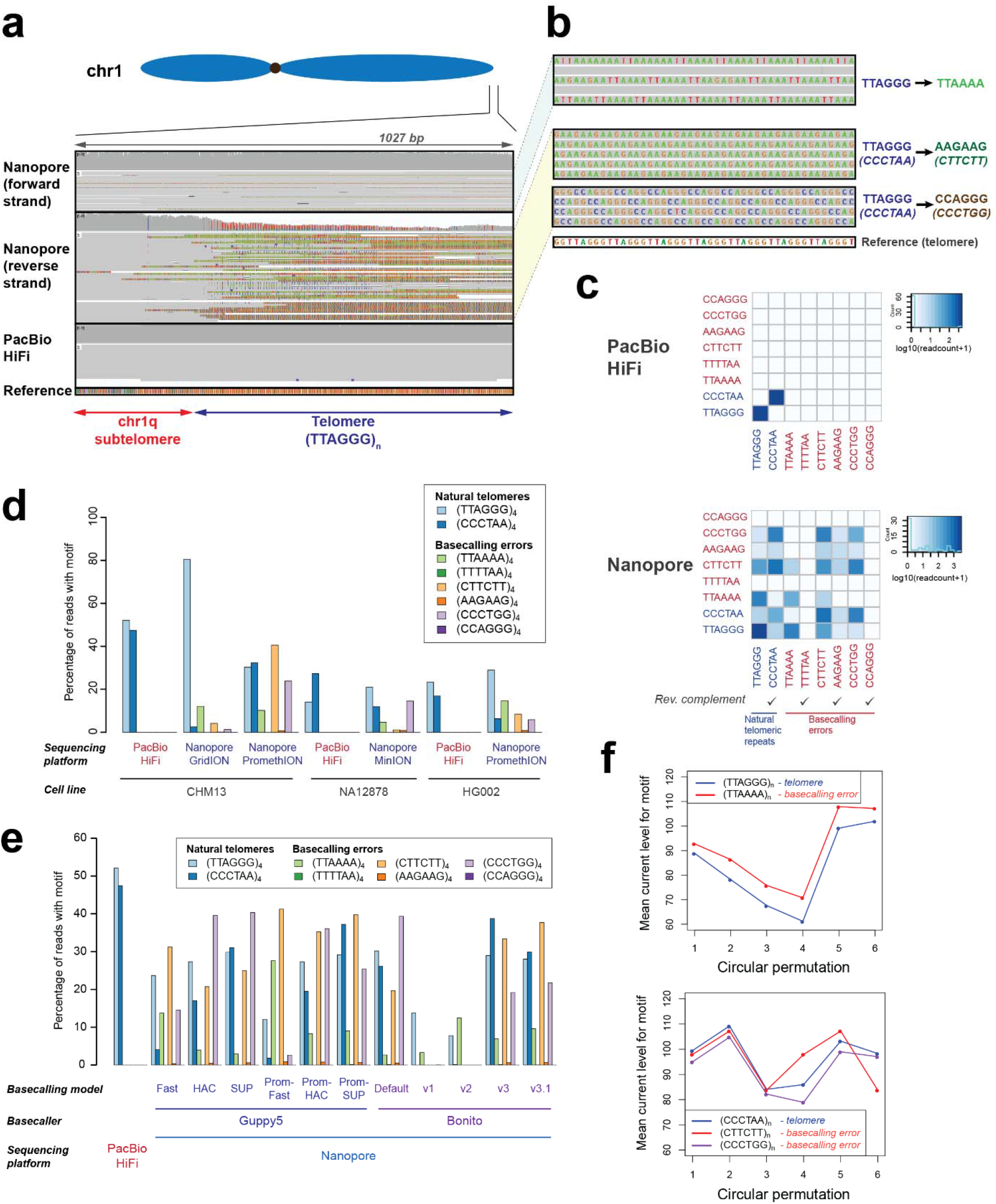
Strand-specific Nanopore basecalling errors are pervasive at telomeres. **(a,b)** IGV screenshot illustrating the three types of basecalling errors found on the forward and reverse strands of telomeres for Nanopore sequencing. (TTAGGG)_n_ on the forward strand of Nanopore sequencing data was basecalled as (TTAAAA)_n_ while (CCCTAA)_n_ on the reverse strand was basecalled as (CTTCTT)_n_ and (CCCTGG)_n_. PacBio HiFi data generated from the same cell line (CHM13) is depicted as a control. Reference genome indicated in the plot corresponds to the chm13 draft genome assembly (v1.0). **(c)** Co-occurrence heatmap illustrating the frequency of co-occurrence of repeats corresponding to natural telomeres, or to basecalling errors in PacBio HiFi and Nanopore long-reads found at chromosomal ends (within 10kb of annotated end of the reference genome). Diagonal of co-occurrence matrix represents counts of long-reads with only a single type of repeats observed. **(d)** Basecalling errors at telomeres are observed across different nanopore datasets and sequencing platforms. **(e)** Basecalling errors at telomeres are observed different nanopore basecallers and basecalling models. Guppy5 and the Bonito basecallers, and different bascalling models for each bascaller, were used to basecall telomeric reads in the CHM13 PromethION dataset (reads that mapped to flanking 10kb regions of the CHM13 reference genome). **(f)** Basecalling errors share similar nanopore current profiles as telomeric repeats. Current profiles for telomeric and basecalling error repeats were plotted based on known mean current profiles for each k-mer (Methods).

Together, our results suggest that telomeric regions are frequently misrepresented as other types of repeats in a strand-specific manner during Nanopore sequencing.

We then assessed if these errors are broadly observed in other studies or are specific to the CHM13 dataset from the Telomere-to-Telomere consortium. To assess this, we examined the previously published NA12878 and HG002 Nanopore genome sequencing datasets [12,13,16]. Remarkably, the same basecalling errors, TTAGGG→TTAAAA, CCCTAA→CTTCTT, and CCCTAA→CCCTGG, were similarly observed at telomeres in these datasets **(Figure 1d, Supplementary Figure 4a)**, suggesting that these basecalling errors at telomeres are broadly observed across multiple studies. Remarkably, between 40-60% of reads at telomeric regions in these three datasets display at least one of these type of basecalling repeat artefacts for the Nanopore sequencing platform **(Supplementary Figure 4b)**, while these errors were not observed in the PacBio HiFi datasets for the same samples **(Supplementary Figure 4b)**. Further, we also partitioned these datasets based on the sequencing platforms used to generate them, and noted that basecalling error repeats are observed across all three nanopore sequencing platforms (MinION, GridION, PromethION) **(Figure 1d, Supplementary Figure 4a)**. Together, these results show that these error repeats extend across nanopore sequencing datasets and sequencing platforms.

We then questioned if these error repeats are unique to specific nanopore basecallers or basecalling models. We extracted reads from chromosomal ends, and re-basecalled ionic current data of these reads using different basecallers and basecalling models. Using the production-ready basecaller Guppy5 (Oxford Nanopore Technologies), and the developmental-phase basecaller Bonito (Oxford Nanopore Technologies), we noticed that these basecalling error repeats can be readily observed across both basecallers **(Figure 1e, Supplementary Figure 5 and 6)**. Further, these error repeats were also observed when different basecalling models were applied **(Figure 1e)**. Significantly, we also observed that the “fast” basecalling mode in Guppy led to almost complete loss of the (CCCTAA)_n_ strand **(Figure 1e, Supplementary Figure 5a)**, while the “HAC” basecalling model enabled both strands to be recovered, highlighting that the basecalling model applied can affect strand-specific recovery of telomeric reads. Together, these results suggest that error repeats are observable across nanopore basecallers, and basecalling models.

To determine the cause for these repeat-calling errors, we then examined the ionic current profiles of these repeats. We thus generated ionic current profiles of these telomeric repeats and these error repeats, induced by the nanopore basecallers, using known mean current values of different 6-mers **(Methods)**. Remarkably, we observed a high degree of similarity between current profiles between telomeric repeats and these basecalling errors **(Figure 1f)**. Specifically, we observed that (TTAGGG)_n_ telomeric repeats had a high degree of similarity with the (TTAAAA)_n_ error repeats generated by the Bonito base-caller (Pearson correlation = 0.9928, Euclidean distance=4.9934) **(Supplementary Figure 7a-c)**. Similarly, (CCCTAA)_n_ current profile also showed high similarity with (CCCTGG)_n_ repeats (Pearson correlation = 0.9783, Euclidean distance = 4.687), and reasonably good similarity with (CTTCTT)_n_ repeats (Pearson correlation = 0.6411, Euclidean distance = 19.384) **(Supplementary Figure 7a-c)**,. Together, these results suggest that similarities in current profiles between repeat sequences are possible causes for repeat-calling errors at telomeric repeats.

We then examined if repeat-calling errors may extend to other repetitive sequences beyond telomeric sequences. To address this, we search for other repeat pairs with similar current profiles that may be susceptible to these repeat-calling errors. We simulated and performed pairwise comparison of current profiles for all 6-mer repeats (n=8,386,560 comparisons) **(Methods)**. Using similar Pearson correlation (≥0.99) and Euclidean distance cutoffs (≤5) as observed for telomeric repeat errors identified in this study **(Supplementary Figure 7a-c)**, we identified a further 2577 pairs of repeats with similar current profiles **(Supplementary Table 1, Supplementary Figure 7d)**. For instance, we found that (TTAGGG)_n_ telomeric repeats also showed high similarities in current profiles with repeats with single-nucleotide substitutions like (TT<u>AA</u>GG)_n_, (TTAG<u>A</u>G)_n_ and (TT<u>G</u>GGG)_n_ **(Supplementary Figure 7d,e)**. Repeat sequences like (GCTGCT)_n_ and (<u>AAC</u>G<u>GC</u>)_n_ that differed drastically at the sequence level, but shared similar current profiles were also observed **(Supplementary Figure 7d,f)**. Further, we also examined the unmappable pool of CHM13 nanopore reads after mapping it to the CHM13 reference assembly. Remarkably, a significant pool of reads with long (GT)_n_ repeats were readily observed **(Supplementary Figure 8)**. Interestingly, (GTGTGT)_n_ repeats were also found to have high similarities in current profiles with (CTCTCT)_n_ repeats **(Supplementary Figure 7d, Supplementary Table 1)**, suggesting that the pool of unmappable (GT)_n_ reads may include (CT)_n_ repeats. Collectively, our results suggests that these basecalling error repeats may be observed at other repetitive regions, beyond telomeres.

To resolve these basecalling errors at telomeres, we then attempted to tune the nanopore basecaller by providing it with more training examples of telomeres **(Figure 2a)**. Notably, model training was performed with a low learning rate to ensure that the majority of the model does not get affected during training while ensuring that minor adjustments in the model can be made to accurately basecall telomeres. Specifically, we tuned the deep neural network model underlying the Bonito basecaller by training it at a low learning rate with ground truth telomeric sequences extracted from the CHM13 reference genome, and current data of the corresponding reads **(Methods)**. As two Nanopore PromethION runs were performed on the CHM13 dataset, we used the data from one run for training (run225) and tuning of the basecaller, and held out the data from the second run (run 226) for evaluation of our tuned basecaller. With this approach, we see a significant improvement in the base-calls of both the telomeres, and sub-telomeric regions on the training data and held out dataset with clearly observable decrease in errors on the chromosomal ends **(Figure 2b, Supplementary Figure 9a-d)**. Together, our results indicate that a nanopore base-caller can be tuned to more accurately base-call telomeric regions by providing additional training examples.

**Figure 2.**
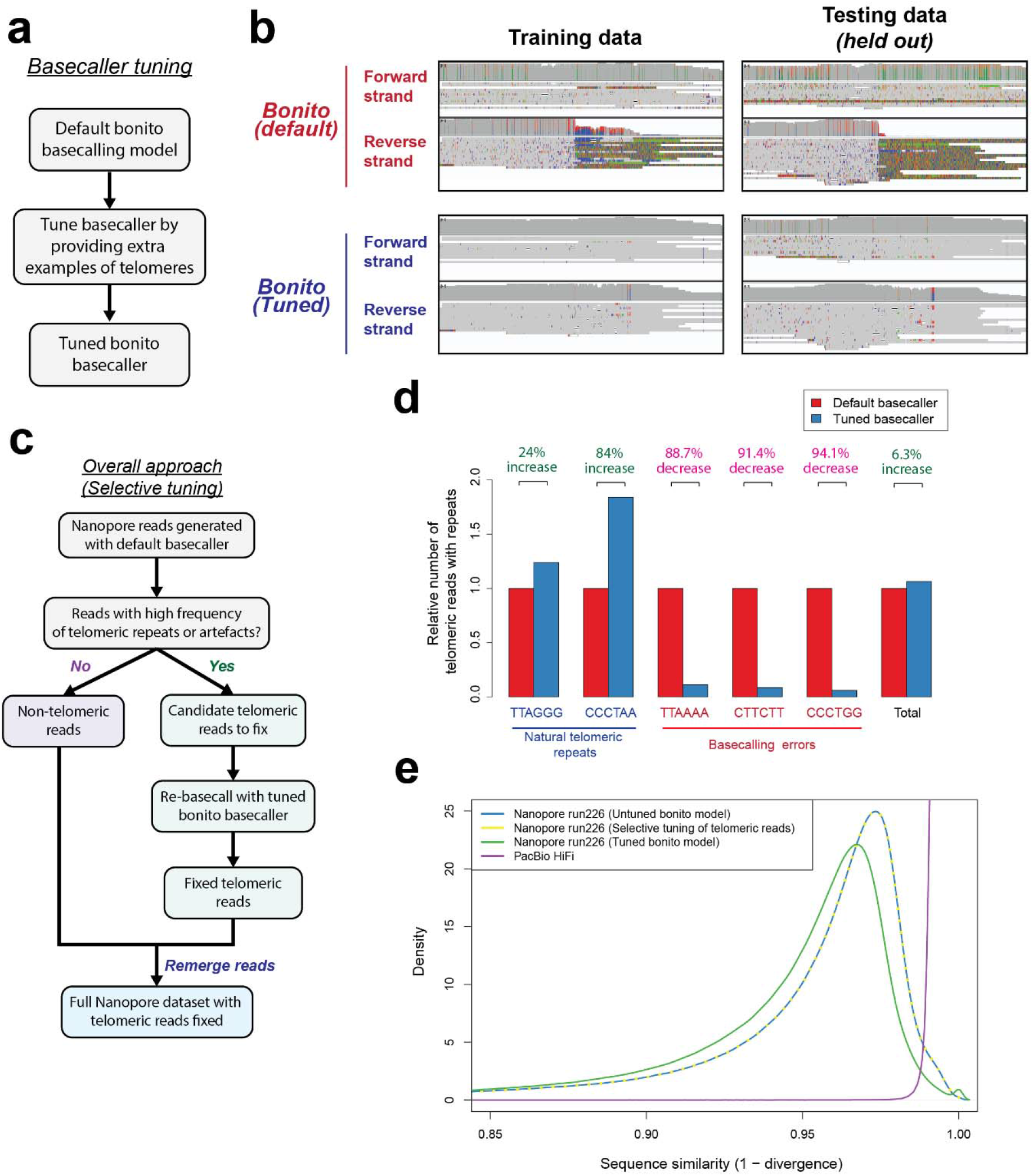
Selective re-basecalling of telomeric reads resolves basecalling errors at telomeres. **(a)** Approach for tuning the bonito basecalling model for improving basecalls at telomeres. **(b)** Tuned bonito basecalling model leads to improvement in basecalls at telomeric regions. IGV screenshots of telomeric region (chr2q) in the CHM13 dataset basecalled using the default bonito basecaller, and the tuned bonito basecalling model is as depicted. **(c)** Overall approach for selecting and fixing telomeric reads in nanopore sequencing datasets. Telomeric reads are selected (Methods), and rebasecalled using the tuned bonito basecalling model. **(d)** The selective tuning approach leads to improved recovery of telomeric reads, and decrease in the number of reads with basecalling artefacts. Evaluation was performed on the held out test dataset (run226). **(e)** The ‘selective basecalling’ approach leads to little detected negative impact on basecalling of other genomic regions. The sequence similarity of all reads to the reference genome for three approaches for basecalling of nanopore reads was evaluated. They are applying the default bonito basecalling model to all reads (untuned bonito model), applying the tuned bonito basecalling model to all reads (tuned bonito model), and applying the tuned bonito basecalling model selectively to telomeric reads only (selective tuning of telomeric reads). The density plot depicts the sequence similarity of each read against the CHM13 reference genome as assessed using minimap2.

As it is computationally more efficient to redo repeat-calling only for the small fraction of problematic telomeric reads rather than all reads, we developed an overall strategy to select these telomeric reads for re-basecalling with the tuned Bonito+telomeres basecaller **(Figure 2c)**. To select telomeric reads for selective re-basecalling, we relied on an observation from the CHM13 reference genome and nanopore sequencing datasets. Specifically, we noticed that telomeric reads which maps to the ends of the CHM13 reference genome tend to show a high frequency of telomeric, or basecalling error repeats as compared to the rest of the genome **(Supplementary Figure 10).** We therefore utilized this observation to separate the non-telomeric reads, from the candidate telomeric reads **(Figure 2c, Methods)**. These telomeric reads were then re-base-called with the tuned Bonito basecaller before being recombined with the pool of non-telomeric reads. Remarkably, with this strategy, we observed a significant improvement in recovery of telomeric reads with (TTAGGG)_n_ and (CCCTAA)_n_ repeats (from 384 to 476 TTAGGG and 373 to 686 CCCTAA reads) **(Figure 2d)**. At the same time, a sharp reduction of these basecalling repeat errors was also observed (151 to 17 TTAAAA reads, 561 to 48 CTTCTT reads, and 337 to 20 CCCTGG reads) **(Figure 2d)**. Together, these results suggests that our “selective tuning” approach for fixing basecalling errors at telomeres can improve recovery of telomeric reads while reducing telomeric basecalling repeat artefacts.

We further evaluated our approach for possible impact on overall basecalling accuracy. While a reduction in global basecalling accuracy was observed (~1-2%) when our tuned basecaller was directly applied to the full dataset, caused likely by miscalling of endogenous (CTTCTT)_n_ genomic repeats as (CCCTAA)_n_, this loss of global basecalling accuracy could be avoided by applying our basecaller to telomeric reads alone. Concordant with this, we did not observe changes in overall basecalling accuracy with our telomere-selective tuning approach **(Figure 2e)**. These results indicate that our telomere-selective tuning approach has negligible impact on basecalling accuracy for the rest of the genome.

## Conclusion

In this study, we showed that basecalling errors can be widely observed at telomeric regions across nanopore datasets, sequencing platforms, basecallers, and basecalling models. We further showed that these strand-specific basecalling errors were likely induced by similarities in current profiles between different repeat types. To resolve these basecalling errors at telomeres, we devised an overall strategy to re-basecall telomeric reads using a tuned nanopore basecaller. More broadly, our study highlights the importance of verifying nanopore basecalls in long, repetitive and poorly defined regions of the genome. For instance, this can be done either with an orthogonal platform, or at a minimum by ensuring nanopore basecalls between opposite strands are concordant. In the future, we anticipate that further improvements in the nanopore basecaller or basecalling model as demonstrated in this study will potentially lead to the reduction or elimination of these basecalling artefacts.

## Methods

### Nanopore and PacBio Datasets

Nanopore and PacBio HiFi datasets for the CHM13 sample were downloaded directly from the telomere-to-telomere consortium (https://github.com/marbl/CHM13)

Nanopore dataset for GM12878 was obtained from the Nanopore WGS consortium (https://github.com/nanopore-wgs-consortium/NA12878/blob/master/Genome.md). PacBio HiFi dataset for GM12878 was obtained from the repository at the SRA database (SRP194450), and downloaded from the following link (https://www.ebi.ac.uk/ena/browser/view/SRR9001768?show=reads)

The HG002 PacBio HiFi and Nanopore datasets were downloaded from the Human Pangenome Reference Consortium (https://github.com/human-pangenomics/HG002_Data_Freeze_v1.0). Specifically, the HG002 Data Freeze (v1.0) recommended downsampled data mix was downloaded. The PacBio HiFi dataset corresponds to ~34X coverage of Sequel II System with Chemistry 2.0. The Nanopore dataset corresponds to 60x coverage of unsheared sequencing from 3 PromethION flow cells from Shafin et al [17].

### Extraction of candidate telomeric reads

Telomeric reads were extracted by mapping all reads to the CHM13 draft genome assembly (v1.0) obtained from the telomere-to-telomere consortium using Minimap2 (version 2.17-r941). Subsequent to that, reads that mapped to within 10 kilobasepairs of the start and end of each autosome and X-chromosome were then extracted using SAMtools (version 1.10).

### Co-occurrence matrix

Candidate PacBio HiFi and Nanopore telomeric reads were first extracted as described above, and then converted into the FASTA format using SAMtools (version 1.10). Subsequent to that, custom Python scripts were used to assess if each of the reads contain at least four consecutive counts of the repeat sequence of interest (e.g. (TTAGGG)_4_). This information is then used to generate a pair-wise correlation matrix as depicted with R in the main text.

### Basecalling of nanopore data with different basecallers and basecalling models

Basecalling of Nanopore data was done using Guppy (Version 4.4.2), Guppy (Version 5.0.16) and Bonito v0.3.5 (commit d8ae5eeb834d4fa05b441dc8f034ee04cb704c69). For Guppy4, four different basecalling models were applied (guppy_dna_r9.4.1_450bps_fast, guppy_dna_r9.4.1_450bps_hac, guppy_dna_r9.4.1_450bps_prom_fast, guppy_dna_r9.4.1_450bps_prom_hac). For Guppy 5, six different basecalling models were applied (dna_r9.4.1_450bps_fast, dna_r9.4.1_450bps_hac, dna_r9.4.1_450bps_sup, dna_r9.4.1_450bps_fast_prom, dna_r9.4.1_450bps_hac_prom, dna_r9.4.1_450bps_sup_prom) For Bonito, the v1, v2, v3, v3.1 and default basecalling models were applied.

### Current profiles for different repeat sequences

The mean current level for different k-mers sequenced by Nanopore sequencing was obtained from the k-mer models published by Oxford Nanopore (https://github.com/nanoporetech/kmer_models/tree/master/r9.4_180mv_450bps_6mer).

Circular permutations of each 6-mer of interest was generated, and their corresponding mean current level extracted from the k-mer models. The current profiles for each of the indicated repeat sequences were then plotted and depicted in the figure.

### Pairwise comparison of all possible k-mers

Current profile for each 6-mer repeat sequence was generated using the published k-mer models as described above. Pairwise comparisons of all possible 6-mer repeat current profiles was then performed (8,386,560 pairs in total). A corresponding (i) Pearson correlation value, (ii) mean-centered Euclidean distance, and (iii) mean current difference for each pair of 6-mer repeat current profiles were then generated. Pairs of repeats with a Pearson correlation value ≥ 0.99 and Euclidean distance ≤ 5 were selected as putative repeat pairs that can be miscalled.

### Tuning of bonito model

The default model from Bonito v0.3.5 (commit d8ae5eeb834d4fa05b441dc8f034ee04cb704c69) was used as the base model for model tuning. The training dataset needed for the training process was generated from the telomeric reads from a PromethION run in the CHM13 dataset (run225). More broadly, we then generate the training dataset by matching the current profiles from the Nanopore run to ground truth sequences that we extracted from the CHM13 draft reference genome assembly (v1.0) using custom written code.

Specifically, these telomeric reads were first basecalled using the initial Bonito basecalling model, and then mapped back to the CHM13 draft reference genome assembly (v1.0). This allowed each telomeric read to be properly assigned to its corresponding chromosomal arm with its sub-telomeric sequence. Nonetheless, as the telomeric region of the same read could not be properly mapped to the telomeric repeats due to the repeat errors, there was difficulty in assigning the nanopore current data to the correct ground truth sequences in the reference genome. As such, the presume length of sequences to extract was estimated using the basecalling repeat error sequences, and the same length of sequences were then extracted from the CHM13 reference genome to serve as ground truth sequences. With this idea and with custom Perl script, we were able to generate a set of ground truth sequences and signals for model tuning. These data were then formatted into the corresponding python objects required by the Bonito basecaller with custom Python scripts. Using the tune function in Bonito and with our prepared training dataset, we were then able to train the basecaller to convergence.

### Selective application of tuned basecaller to telomeric reads

We applied our tuned basecaller by first extracting candidate telomeric reads for re-basecalling. This was done by enumerating the total 3-mer telomeric (TTAGGG, CCCTAA) and repeat artefact count (TTAAAA, CTTCTT, CCCTGG) on each read. Reads with at least 10 total counts of these repeats were isolated and their readnames noted. These reads were then excluded from the total pool of reads via their readnames, and basecalled separately using our tuned basecaller using the fast5 data of these reads. Following basecalling with the tuned basecaller, these reads were then recombined with the main pool of reads.

## Abbreviations

PacBio: Pacific Biosciences
SMRT: Single Molecule Real Time

## Author information

### Author contributions

K.T.T. and M.S. identified issues with Nanopore sequencing of telomeres, and discovered basecalling errors at telomeric regions. K.T.T. evaluated basecalling errors in Nanopore sequencing datasets, and designed the overall approach for correcting basecalling errors at telomeric regions with inputs from H.L. and M.M. K.T.T. wrote the initial draft of the manuscript with inputs from H.L. and M.M. M.M. and H.L. jointly supervised the work. All authors read, revised, and approved the submission of the manuscript.

## Declarations

### Ethics approval and consent to participate

Not applicable.

### Consent for publication

Not applicable.

### Availability of data and materials

Source code to apply and retrain the bonito bascalling model for telomeric region can be found at the following link: https://github.com/ktan8/nanopore_telomere_basecall/.

The tuned bonito basecalling model can be downloaded from https://zenodo.org/api/files/86cb9586-300f-493d-b9c4-0ab2f2848e3c/chm13_nanopore_trained_run225.zip. A comprehensive version of Supplementary Table 1 with all possible pairs of k-mers can be found at https://zenodo.org/api/files/86cb9586-300f-493d-b9c4-0ab2f2848e3c/all_comparisions.similar_profile.txt.zip.

## Acknowledgements

We would like to thank all members of the H.L. and M.M. labs for helpful comments and discussions. We would also like to thank the Telomere-to-Telomere consortium for generating the CHM13 datasets used in this study.

---

**Supplementary Figure 1.**
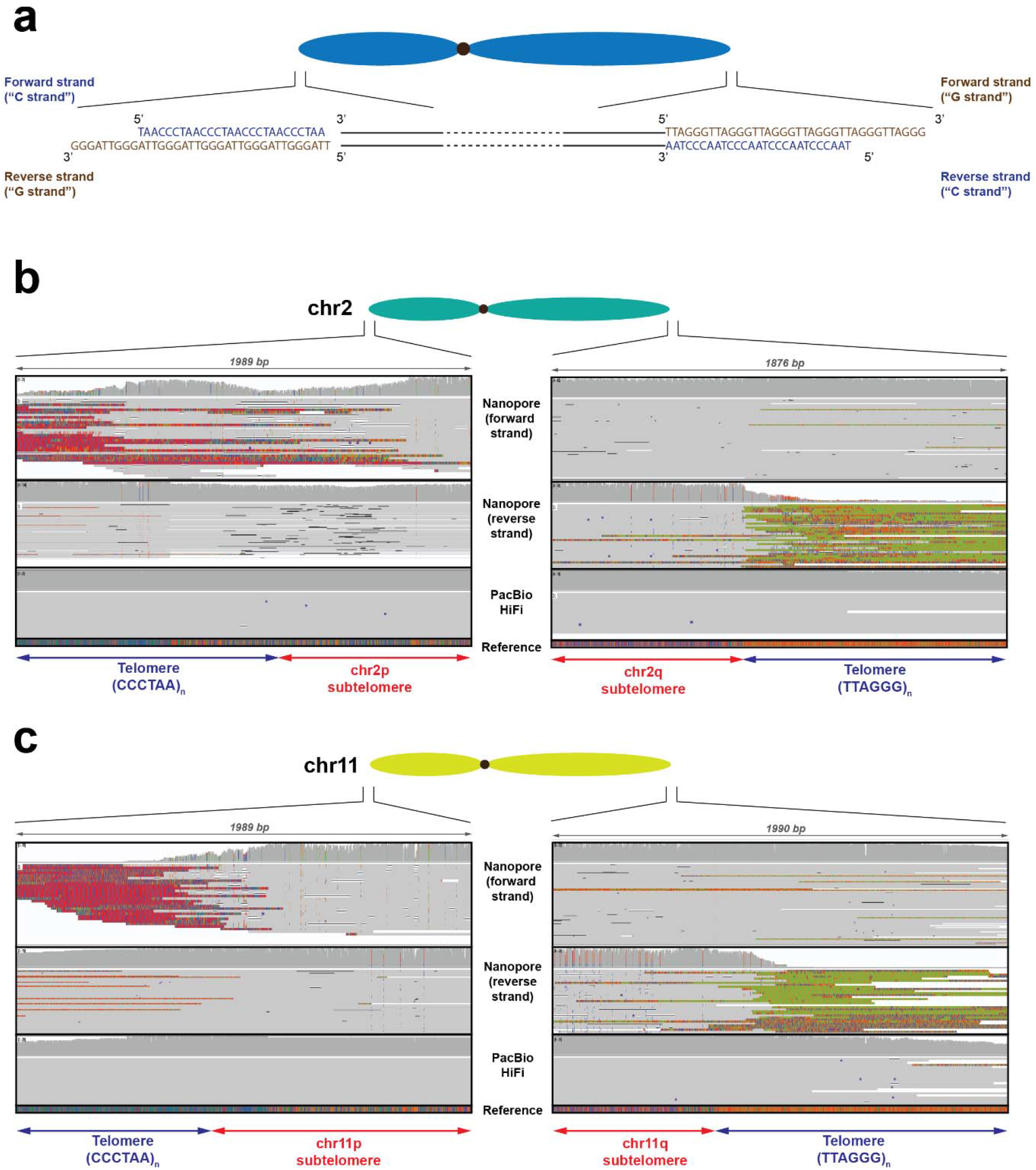
Additional screenshots of basecalling repeat errors found on different chromosomal arms. **(a)** Schematic depicting sequence and orientation of telomeric repeat sequences on the p-arms (arm on the left in the schematic) and q-arms (arm on the right of the schematic) of a chromosome. Note that the forward strand for the arm on the left, and reverse strand for the arm on the right are “C-rich strands” and characterized by (CCCTAA)_n_ repeats in a 5’-to-3’ direction. Also note that the reverse strand for the arm on the left, and forward strand for the arm on the right are “G-rich strands” and characterized (TTAGGG)_n_ repeats in a 5’-to-3’ direction. **(b-c) S**creenshots depicting additional representative examples of chromosomal arms with basecalling error repeats. These are **(b)** chromosome 2 and **(c)** chromosome 11. Screenshots were extracted from the Integrative Genomics Viewer for the CHM13 long-read dataset mapped against the CHM13 reference genome. Related to Figure 1a.

**Supplementary Figure 2.**
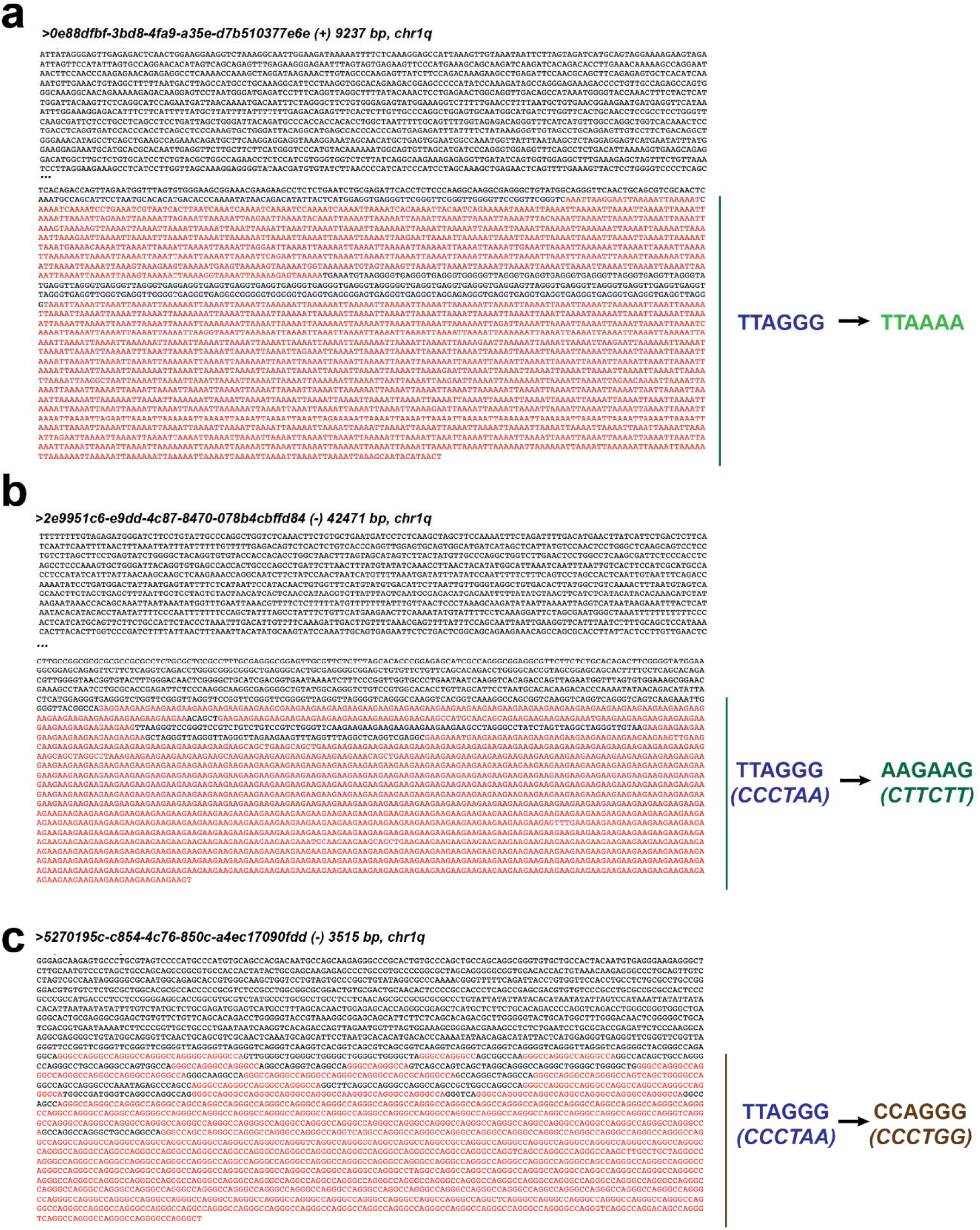
Examples of long-reads with three types of basecalling error repeats found at telomeres. **(a-c)** Sequences and readnames of representative long-reads with the three reported types of basecalling error repeats are as depicted. The region with the basecalling error repeats is highlighted in red. The three type of basecalling errors found on each long read are **(a)** (TTAGGG)_n_ to (TTAAAA)_n_, **(b)** (CCCTAA)_n_ to (CTTCTT)_n_ and **(c)** (CCCTAA)_n_ to (CCCTGG)_n_. Note that **(b)** and **(c)** represents the reverse complementary sequence of the actual nanopore long-read sequence. Also note that the repeats were found on the end of each read as expected given that telomeric repeats are typically found on the end of the chromosomes.

**Supplementary Figure 3.**
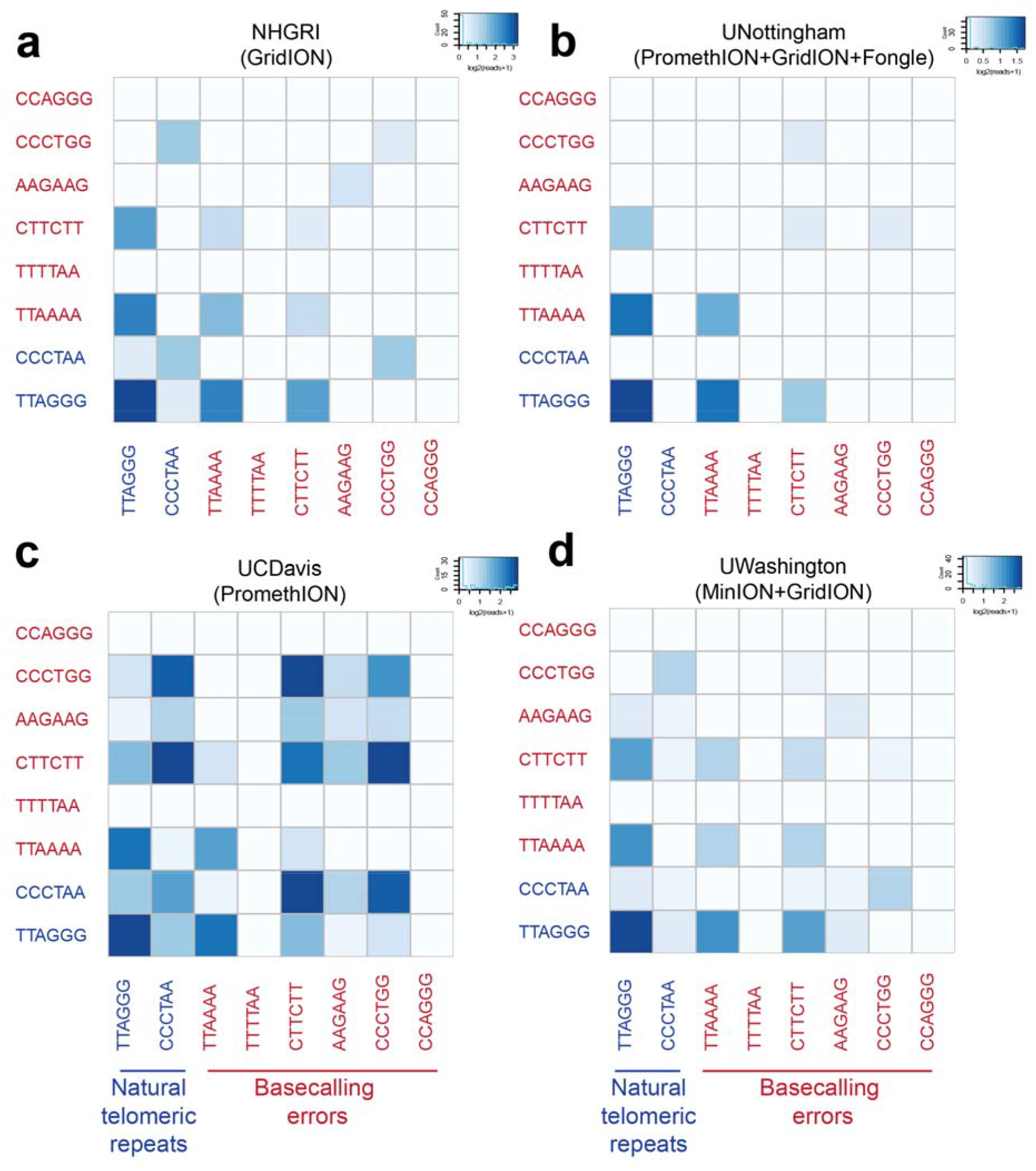
Co-occurrence heatmap illustrating the frequency of co-occurrence of telomeric repeats and basecalling errors for the CHM13 Nanopore dataset generated at different sites. These are **(a)** National Human Genome Research Institute (NHGRI), **(b)** University of Nottingham (UNottingham), **(c)** University of California, Davis (UCDavis) and **(d)** University of Washington (UWashington). The sequencing platforms used for sequencing at each of the sites are also as indicated. This figure is related to Figure 1b.

**Supplementary Figure 4.**
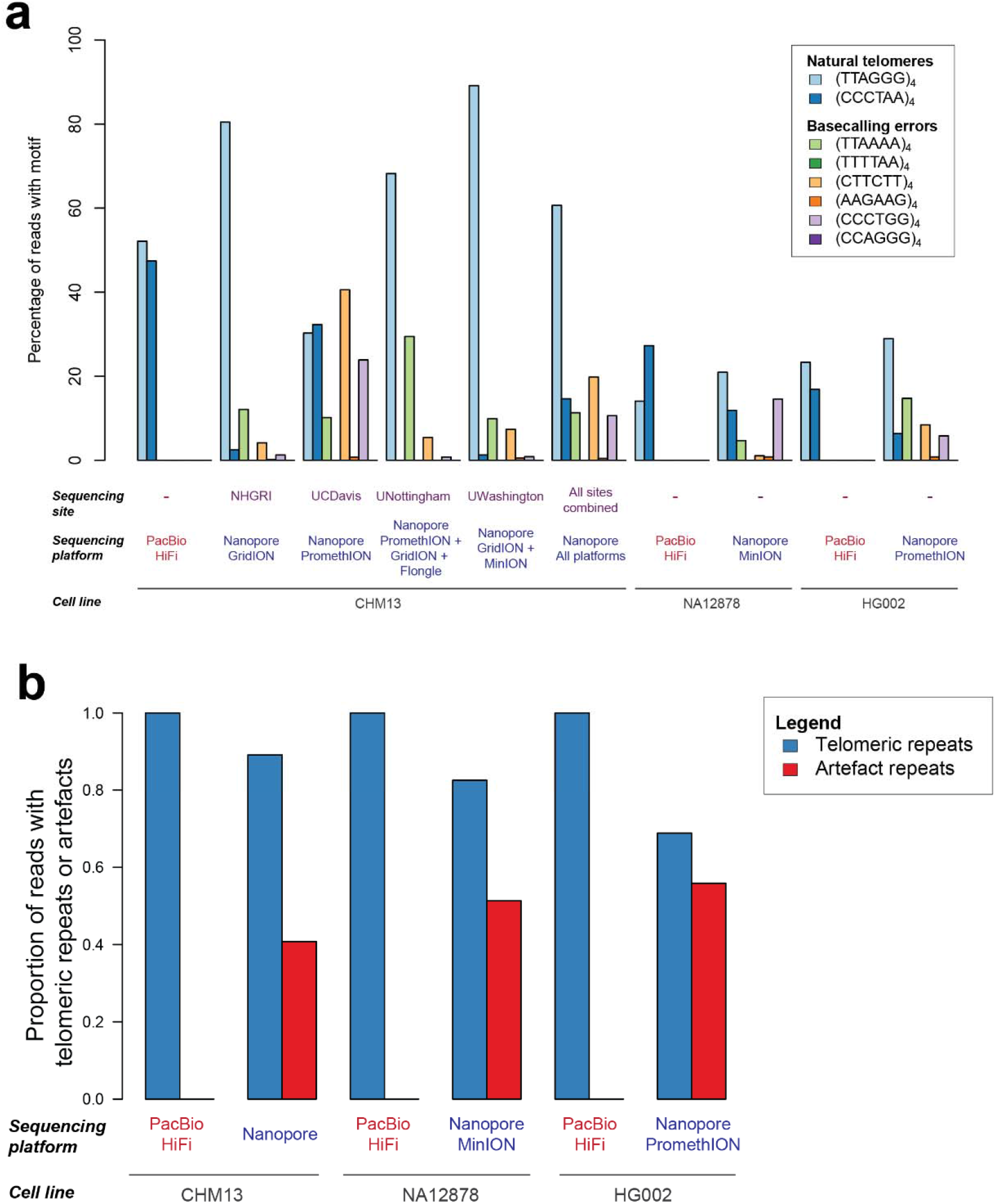
Frequency of telomeric repeat errors in different Nanopore sequencing dataset and sequencing platforms. **(a)** Frequency of basecalling error repeats on three different cell lines generated by different Nanopore sequencing platforms. This figure is an extension Figure 1d. **(b)** Aggregated fraction of basecalling error repeats for different cell lines and datasets.

**Supplementary Figure 5.**
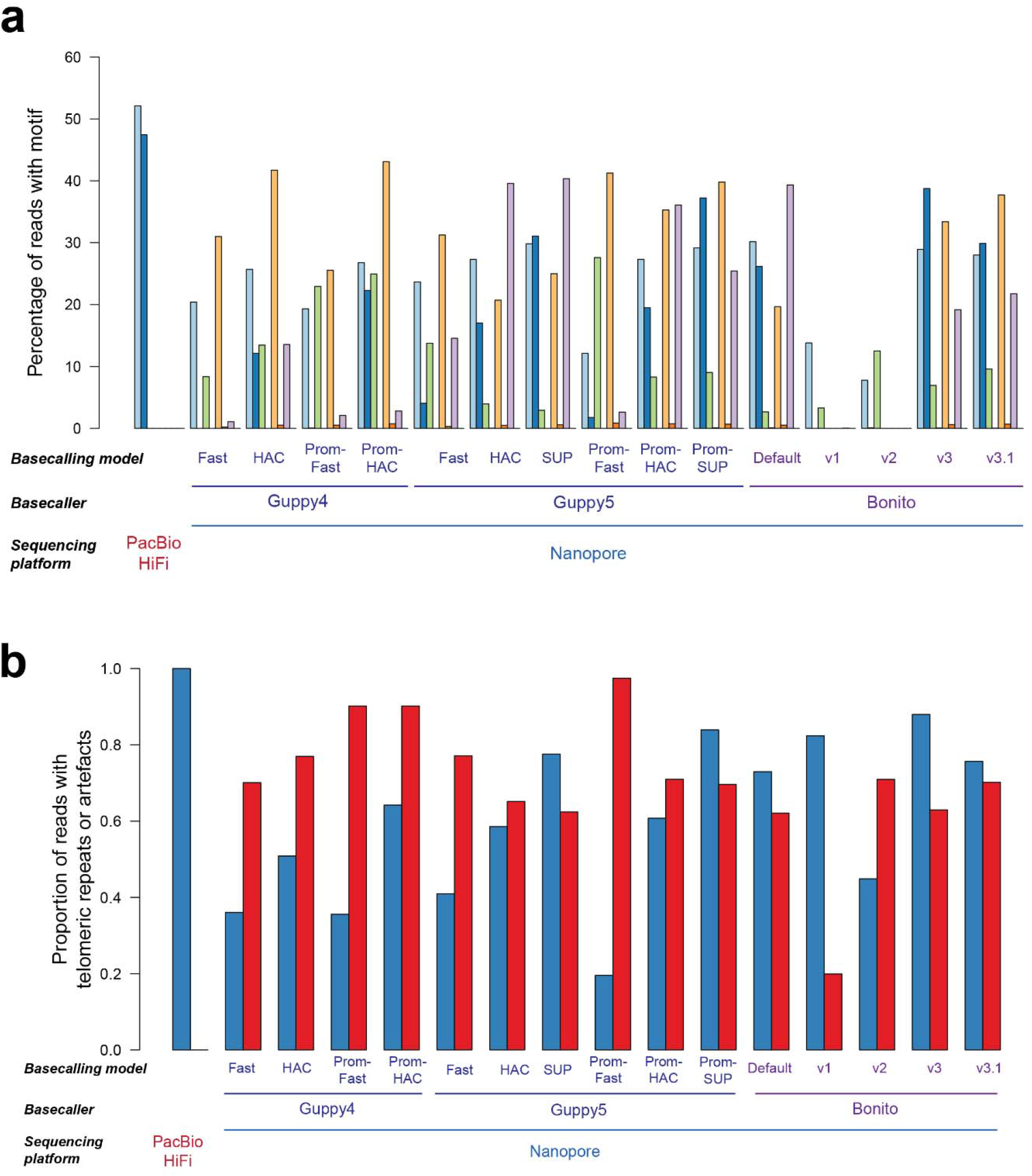
Frequency of telomeric repeat errors in different Nanopore basecallers. **(a)** Frequency of basecalling error repeats for different basecallers (Guppy4, Guppy5 and Bonito) and basecalling models. This figure is an extension of Figure 1e. **(b)** Aggregated fraction of basecalling error repeats for different basecallers and basecalling models.

**Supplementary Figure 6.**
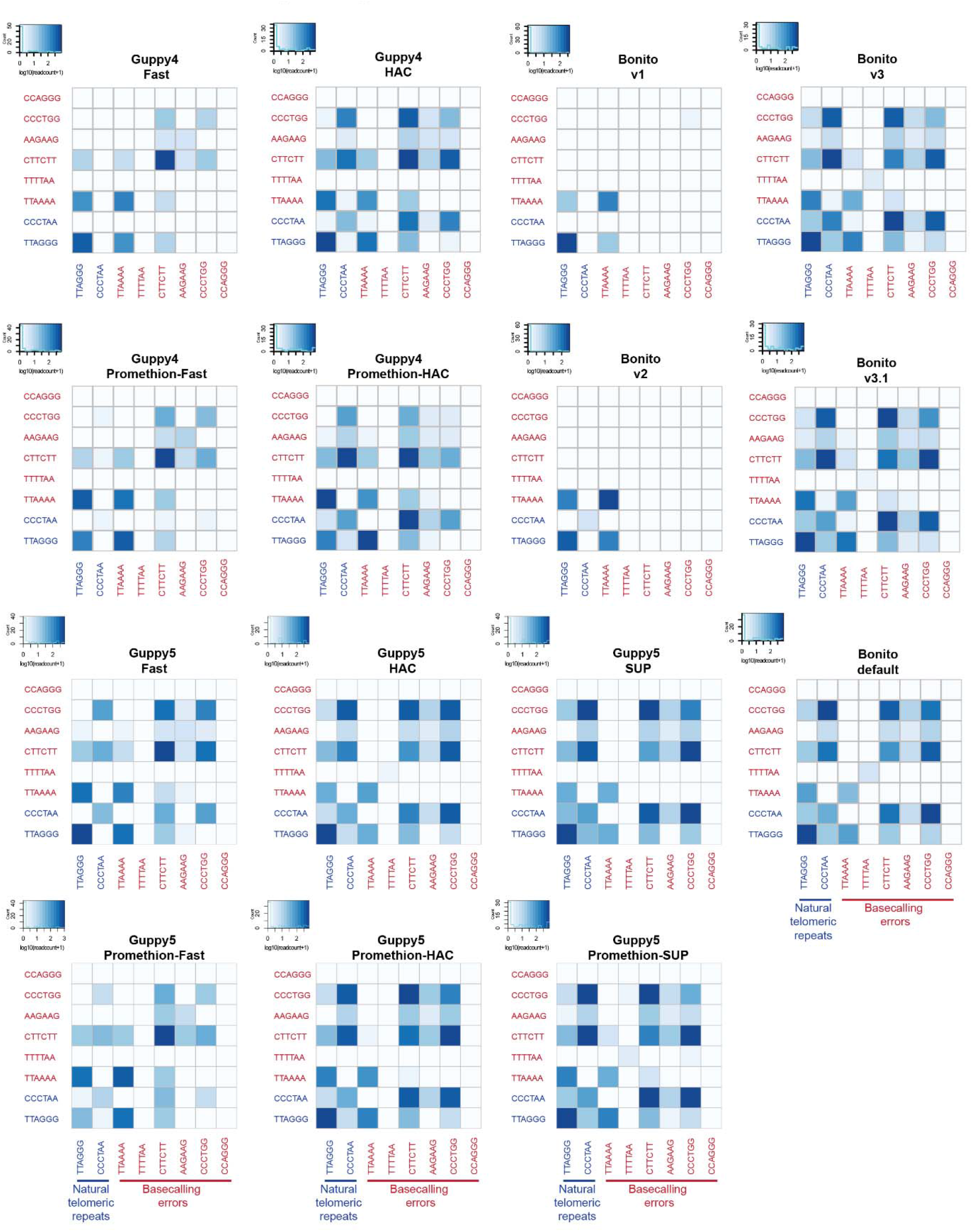
Co-occurrence heatmap for different Nanopore basecalling models. Different nanopore basecallers and basecalling models were applied to the CHM13 Nanopore promethion datasets. The frequency of telomeric repeats and basecalling artefacts observed on reads obtained are as depicted.

**Supplementary Figure 7.**
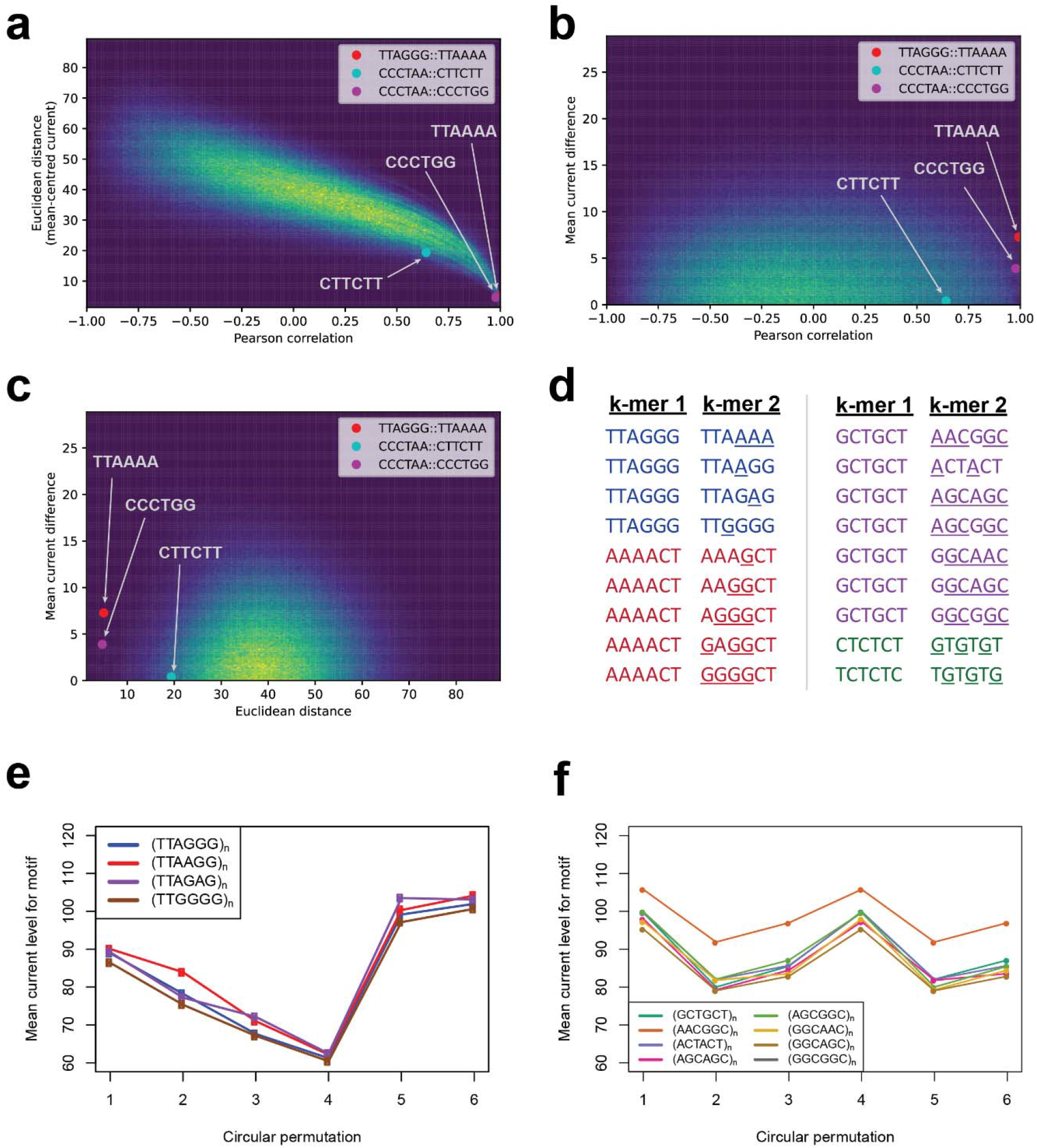
Similarities between current profiles for all possible pairs of 6-mer repeats. **(a-c)** Heatmaps depicting the Euclidean distances, Pearson correlation, and mean current differences between current profiles between all possible 6-mer repeat sequences. These are depicted as pairwise plots for **(a)** the Euclidean distances vs. the Pearson correlation, **(b)** the mean current difference vs. the Pearson correlation, and **(c)** the mean current difference vs. the Euclidean distance. The pairwise comparisons between the telomeric repeats and the observed basecalling repeat artifacts are also highlighted in the plots. **(d)** Example pairs of k-mer repeats with similar current profiles are as indicated. The nucleotides in k-mer 2 that differs from k-mer 1 is underlined to highlight the nucleotides that differ between the two types of repeats. **(e-f)** Current profiles for repeats which were predicted to be highly similar to each other. These are depicted for **(e)** TTAGGG telomeric repeats and telomere-like repeat sequences and (**f)** GCTGCT repeat sequences that were highlighted in purple in Supplementary Figure 7d.

**Supplementary Figure 8.**
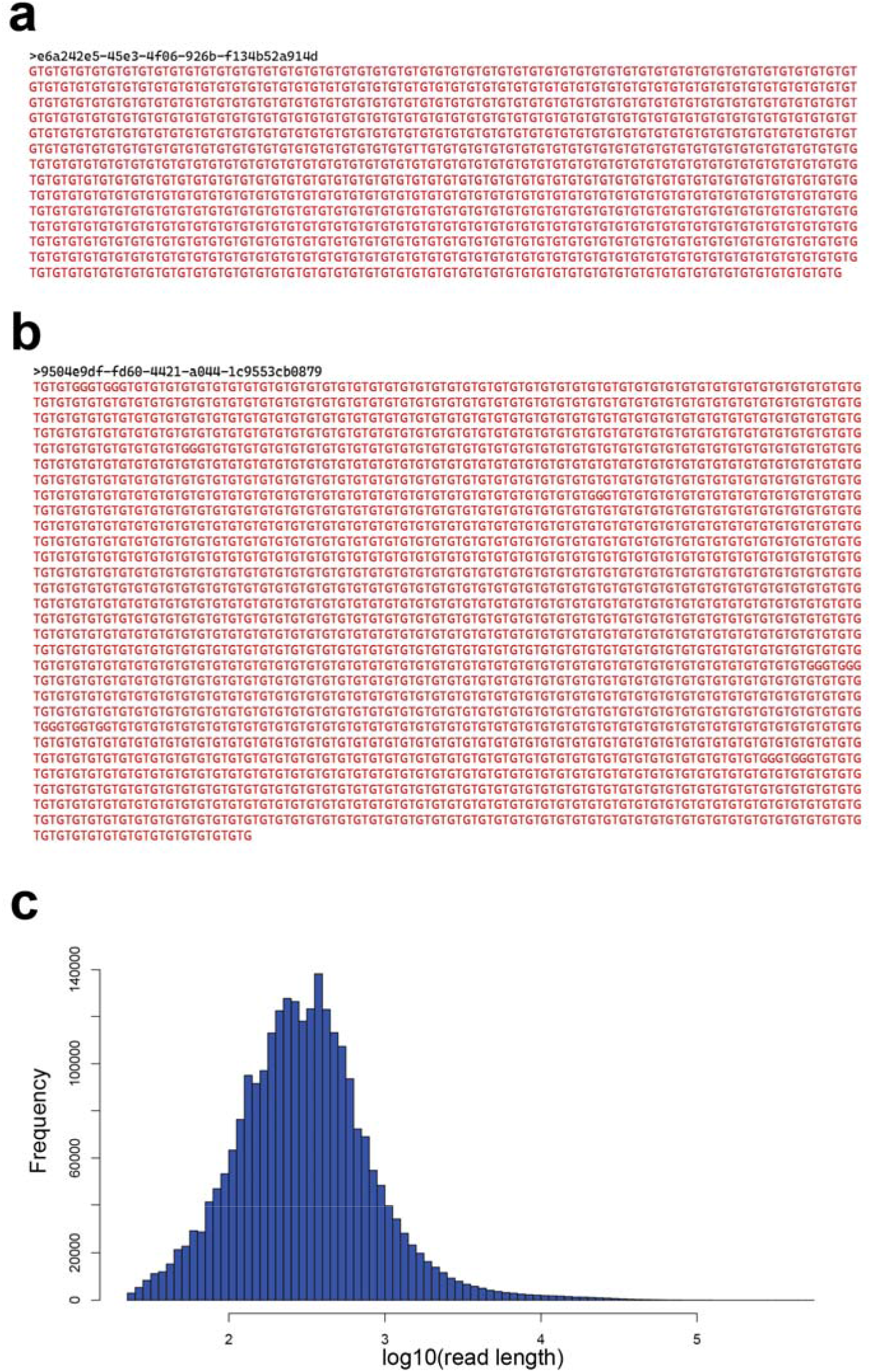
Example of reads with (GT)_n_ repeat sequences in the CHM13 dataset. **(a-b)** Two representative reads from the CHM13 nanopore sequencing dataset with (GT)_n_ repeat sequences. **(c)** Read length distribution of unmappable (GT)_n_ repeats (number of repeats ≥12) in the CHM13 nanopore dataset.

**Supplementary Figure 9.**
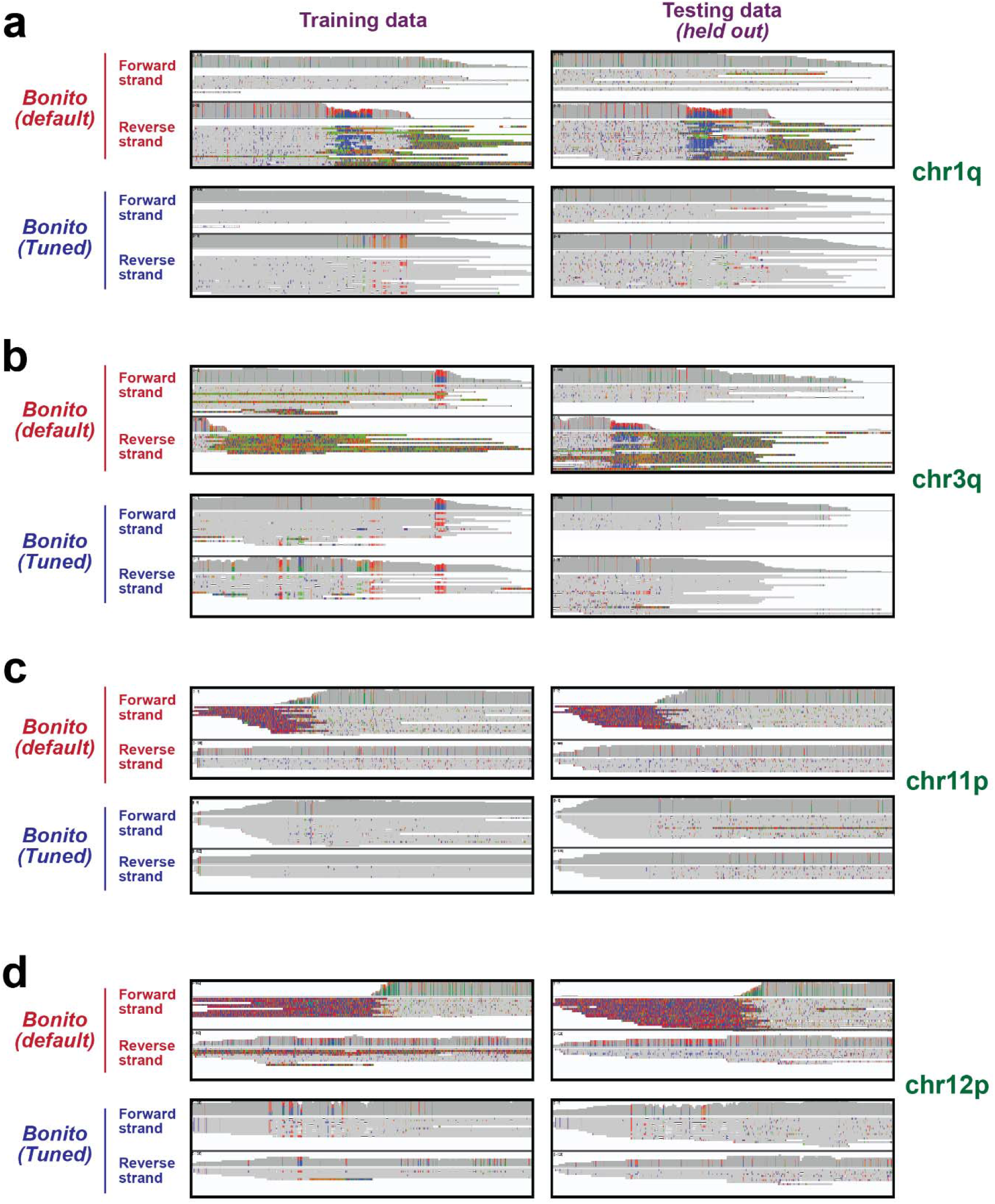
Additional examples for the performance of the tuned bonito basecaller on telomeres on other chromosomal arms. The tuned model was applied to the training dataset used for model training, and on an additional held out test dataset that was not used during model training. IGV screenshots of the default and tuned bonito basecaller on the training and testing dataset for the chromosomal arms **(a)** chr1q, **(b)** chr3q, **(c)** chr11p and **(d)** chr12p are as depicted. Related to Figure 2b.

**Supplementary Figure 10.**
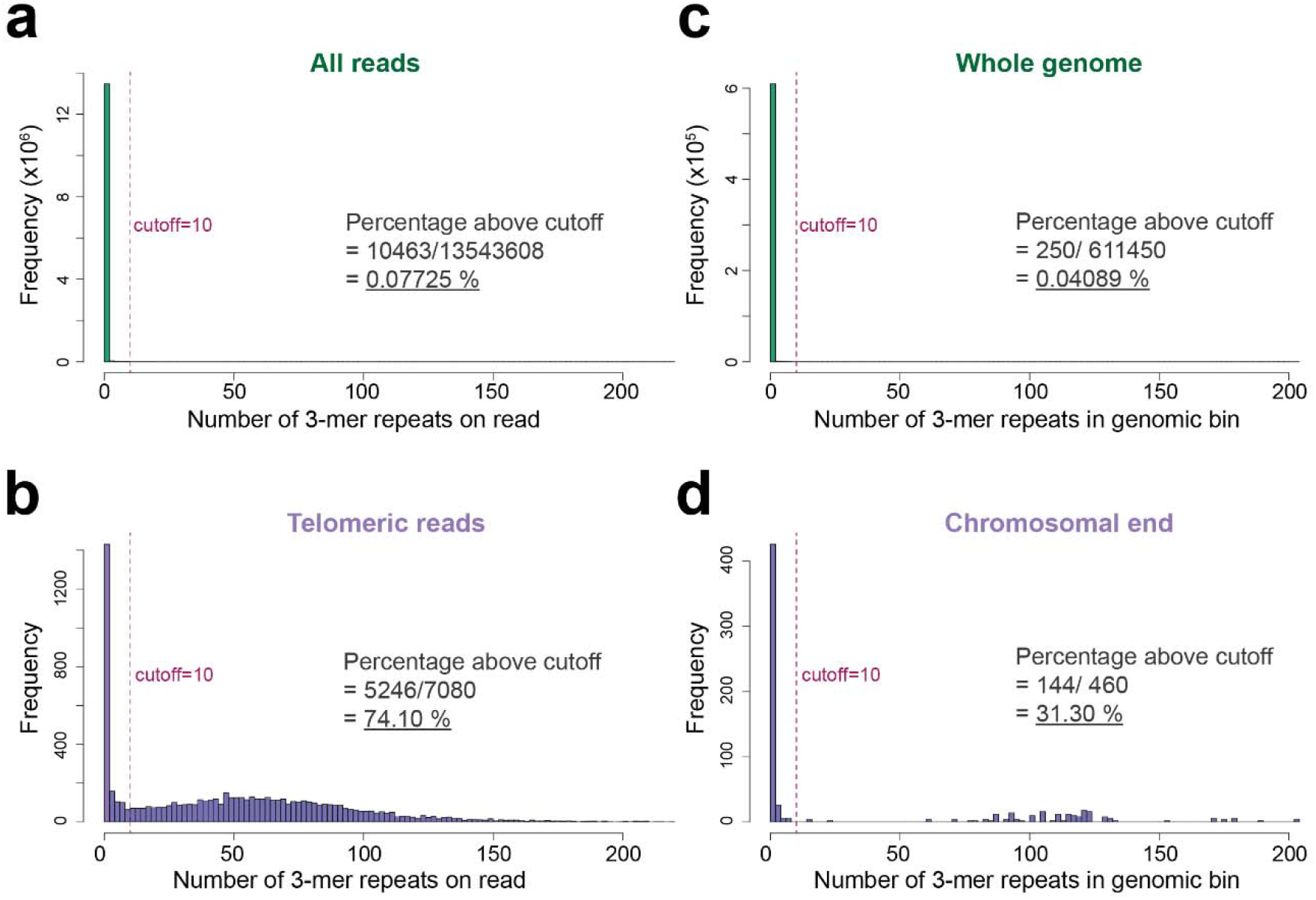
Histograms depicting the frequencies of 3-mer repeats on reads at telomeres and on reads found at the rest of the genome in the CHM13 dataset. **(a-b)** The sum of 3-mer telomeric repeats [(TTAGGG)_3_ (CCCTAA)_3_] and basecalling error repeats [(TTAAAA)_3_, (TTTTAA)_3_, (CTTCTT)_3_, (AAGAAG)_3_, (CCCTGG)_3_, (CCAGGG)_3_] on **(a-b)** each long-read or **(c-d)** genomic bin are as depicted on the x-axis of each histogram. The histograms represent the frequency of these repeats on **(a)** all long-reads in the CHM13 dataset, **(b)** telomeric reads in the CHM13 dataset, **(c)** 20 kb genomic bins with 10 kb moving window for the full CHM13 reference genome, **(d) a**nd for the 10 genomics bins on each chromosomal end of the CHM13 genome.

## Supplementary Tables

**Supplementary Table 1 List of k-mers with high similarities in current profiles.**

The pearson correlation, Euclidean distance, and mean current difference between each pair of k-mer is as presented in the table.

